# Isoform cell type specificity in the mouse primary motor cortex

**DOI:** 10.1101/2020.03.05.977991

**Authors:** A. Sina Booeshaghi, Zizhen Yao, Cindy van Velthoven, Kimberly Smith, Bosiljka Tasic, Hongkui Zeng, Lior Pachter

## Abstract

Full-length SMART-Seq single-cell RNA-seq can be used to measure gene expression at isoform resolution, making possible the identification of isoform markers for cell types and for an isoform atlas. In a comprehensive analysis of 6,160 mouse primary motor cortex cells assayed with SMART-Seq, we find numerous examples of isoform specificity in cell types, including isoform shifts between cell types that are masked in gene-level analysis. These findings can be used to refine spatial gene expression information to isoform resolution. Our results highlight the utility of full-length single-cell RNA-seq when used in conjunction with other single-cell RNA-seq technologies.

## Introduction

Transcriptional and post-transcriptional control of individual isoforms of genes is crucial for neuronal differentiation^1–5^, and isoforms of genes have been shown to be specific to cell types in mouse and human brains^6–11^. It is therefore not surprising that dysregulation of splicing has been shown to be associated with several neurodevelopmental and neuropsychiatric diseases^3,12,13^. As such, it is of interest to study gene expression in the brain at single-cell and isoform resolution.

Nevertheless, current single-cell studies aiming to characterize cell types in the brain via single-cell RNA-seq (scRNA-seq) have relied mostly on gene-level analysis. This is, in part, due to the nature of the data produced by the highest throughput single-cell methods. Popular high-throughput assays such as Drop-seq, 10x Genomics’ Chromium, and inDrops, produce 3’-end reads which are, in initial pre-processing, collated by gene to produce per-cell gene counts. The SMART-Seq scRNA-seq method introduced in 2012^14^ is a full-length scRNA-seq method, allowing for quantification of individual isoforms of genes with the expectation-maximization algorithm^15^. However, such increased resolution comes at the cost of throughput; SMART-Seq requires cells to be deposited in wells, thus limiting the throughput of the assay. In addition, SMART-Seq requires more sequencing per cell.

These tradeoffs are evident in scRNA-seq data from the primary motor cortex (MOp) produced by the BRAIN Initiative Cell Census Network (BICCN). An analysis of 6,160 (filtered) SMART-Seq v4 cells and 90,031 (filtered) 10x Genomics Chromium (10xv3) cells (Figure 1a,b and Extended Data Fig. 1) shows that while 10xv3 and SMART-Seq are equivalent in defining broad classes of cell types, 3’-end technology that can assay more cells can identify some rare cell types that are missed at lower cell coverage (Extended Data Fig. 2a). Overall 125 clusters with gene markers could be identified in the 10xv3 data but not in the SMART-Seq data while only 40 clusters with gene markers could be identified in the SMART-Seq data and not the 10xv3 data, and this differential is consistent with prior comparisons of the technologies^16^. However, while SMART-Seq has lower throughput than some other technologies, it has a significant advantage: by virtue of probing transcripts across their full length, SMART-Seq makes possible isoform quantification and the detection of isoform markers for cell types that cannot be detected with 3’-end technologies (Extended Data Figs. 2b,c). Moreover, SMART-Seq has higher sensitivity than many other methods, which can make possible refined cell type classification.

**Figure 1:**
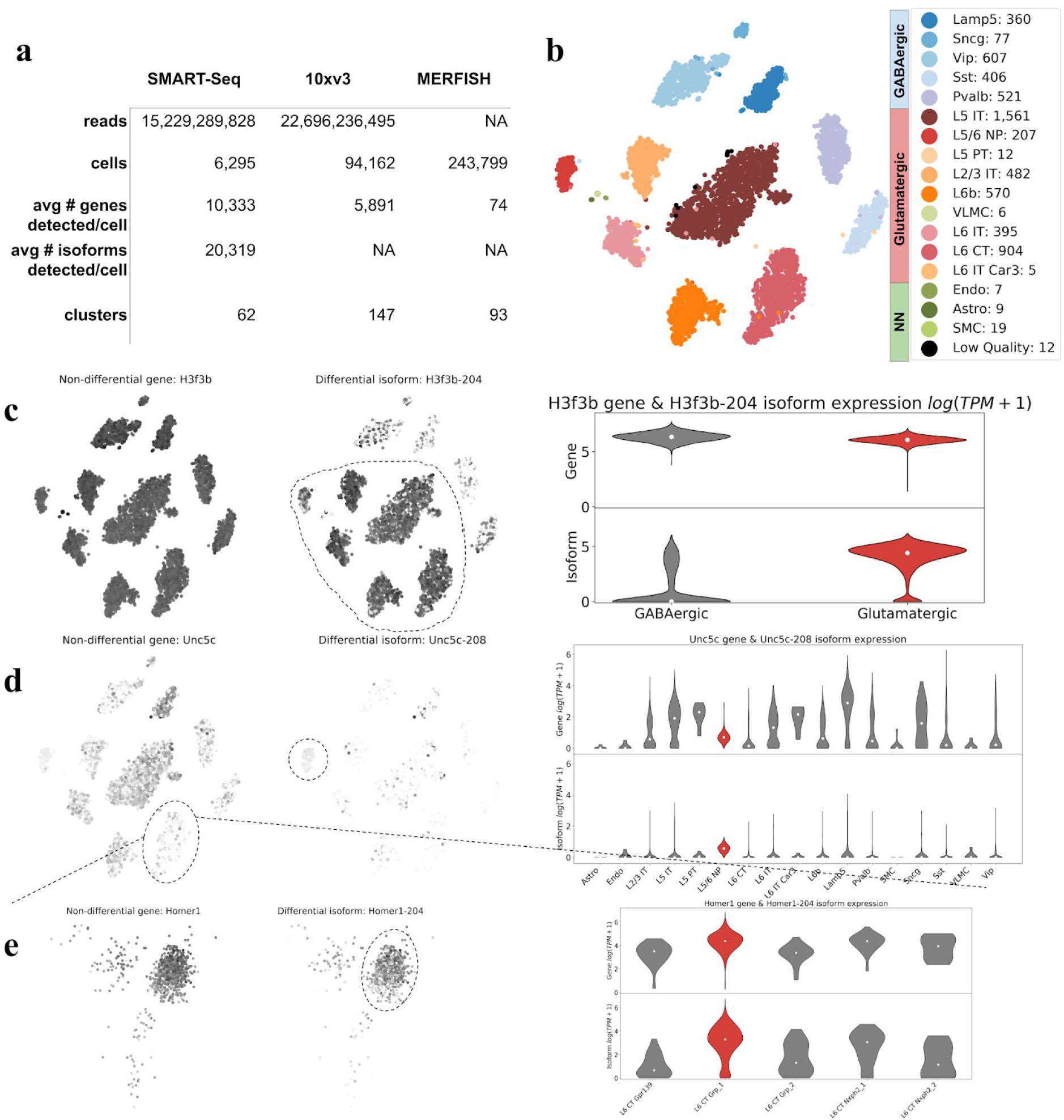
Isoform specificity in the absence of gene specificity. a) Overview of data analyzed. b) A t-SNE map of 10 neighborhood components of 6,160 SMART-Seq cells colored according to cell type. c) The H3f3b gene abundance distribution across cells (left), H3f3b-204 isoform distribution across cells (middle), and violin plots of the gene and isoform distributions. d) Example of a gene with an isoform specific to the L5/6 NP subclass. The Unc5c-208 isoform is highly expressed in L5/6 NP relative to the other subclasses. e) Example of a gene with an isoform specific to the Grp_1 cluster of the L6 CT subclass. The Homer1-204 isoform is highly expressed in L6 CT Grp_1 relative to the other subtypes of L6 CT. [<u>Code a</u>, <u>Code b</u>, <u>Code c</u>, <u>Code d</u>]

By leveraging the isoform resolution of SMART-Seq, we are able to identify isoform-specific markers for dozens of cell types characterized by the BICCN consortium^17^, and find isoform shifts between cell types that are masked in standard analyses. In addition to revealing extensive isoform diversity and cell type specificity in the MOp, our work highlights the complementary value of full-length scRNA-seq to droplet based and spatial transcriptomic methods. Our methods are open-source, reproducible, easy-to-use and constitute an effective workflow for leveraging full-length scRNA-seq data.

## Results

### Isoforms markers for cell types

To identify isoform markers of cell types, we first sought to visualize our SMART-Seq data using gene derived cluster labels from the BICCN analysis^17^. Rather than layering cluster labels on cells mapped to 2-D with an unsupervised dimensionality reduction technique such as t-SNE^18^ or UMAP^19^, we projected cells with neighborhood component analysis (NCA). NCA takes as input not just a collection of cells with their associated abundances, but also cluster labels for those cells, and seeks to find a projection that minimizes leave-one-out k-nearest neighbor error^20^. Thus, while the method outputs a dimension reducing linear map much like principal component analysis (PCA), it takes advantage of cluster labels to find a biologically relevant representation. Visualization with this approach produces meaningful representation of the global structure of the data (Figure 1b), without overfitting (Extended Data Fig. 3a). Moreover, t-SNE applied to PCA (Extended Data Fig. 3b) scrambles the proximity of glutamatergic and GABAergic cell types, while t-SNE of NCA appears to respect global structure of the cells. While UMAP applied to PCA of the data (Extended Data Fig. 3c) appears to be better than t-SNE in terms of preserving global structure, it still does not separate out the cell types as well as NCA (Extended Data Fig. 3d).

Next, motivated by the discovery of genes exhibiting differential exon usage between glutamatergic and GABAergic neurons in the primary visual cortex^11^, we performed a differential analysis between these two classes of neurons. We searched for significant shifts in isoform abundances in genes whose expression was stable across cell types (for details see Methods). We discovered 312 such isoform markers belonging to 260 genes (Supplementary Table 1, [<u>Code</u>]). Figure 1c shows an example of such an isoform from the H3 histone family 3b (H3f3b) gene. The gene has an isoform that is highly expressed in glutamatergic neurons, but the gene undergoes an isoform shift in GABAergic neurons where the expression of the H3f3b-204 isoform is much lower. A gene-level analysis is blind to this isoform shift (top panel, right).

We hypothesized that there exist genes exhibiting cell type isoform specificity at all levels of the MOp cell ontology. However, detection of such genes and their associated isoforms requires meaningful cell type assignments and accurate isoform quantifications. To assess the reliability of the SMART-Seq clusters produced by the BICCN^21^, we examined the correlation in gene expression by cluster with another single-cell RNA-seq technology, the 10xv3 3’-end assay. 90,031 10xv3 cells, also derived from the MOp, were clustered using the same method as the SMART-Seq cells (see Methods). We found high correlation of gene expression between the two assays at the subclass and cluster levels (Extended Data Fig. 4). However, we noted a low correlation in one case, the L5 IT subclass. The low correlation was also observed in a comparison between SMART-Seq and MERFISH gene expression data (Extended Data Fig. 5a), and 10xv3 and MERFISH data (Extended Data Fig. 5b). We hypothesize that this low correlation stems from distinct cell types being clustered together (Extended Data Fig. 5c). To avoid bias, we decided not to search for markers for the L5 IT subclass.

To validate the SMART-Seq isoform quantifications we examined correlations between SMART-Seq and 10xv3 for isoforms containing some unique 3’ UTR sequence. This allowed for a validation of isoform quantifications with a different technology (see Methods). To extract isoform quantifications from 10xv3 data in cases where there was a unique 3’ sequence, we relied on transcript compatibility counts^22^ produced by pseudoalignment with kallisto^23^. We were able to validate the SMART-Seq isoform shift predictions at both the subclass and cluster levels (Extended Data Fig. 6). The isoform abundance correlations are slightly lower than those of gene abundance estimates (Extended Data Fig. 4), but sufficiently accurate to identify significant isoform shifts, consistent with benchmarks showing that isoforms can be quantified accurately from full-length bulk RNA-seq^24^. This is surprisingly accurate considering the underlying differences in the 10xv3 and SMART-Seq technologies.

Having validated the cluster assignments and isoform abundance estimates, we tested for isoform switches for 17 cell subclasses (example in Figure 1d), and then for 54 distinct clusters (example in Figure 1e); see Methods. At the higher level of 17 cell subclasses, we found a total of 913 genes exhibiting isoform shifts among the 18 cell subclasses despite constant gene abundance (Supplementary Table 2, [<u>Code</u>]). We found 40 genes exhibiting isoform shifts among clusters within the L6 CT subclass despite constant gene abundance (Supplementary Table 3, [<u>Code</u>]).

Along with isoforms detectable as differential between cell types without change in gene abundance, we identified isoform markers for the classes, subclasses, and clusters in the MOp ontology that are differential regardless of gene expression. We found 3,911 isoforms belonging to 3,120 genes that are specific to the glutamatergic and GABAergic classes (Figure 2, Supplementary Table 4, [<u>Code</u>]), 2,480 isoforms belonging to 2,146 genes exhibiting isoform shifts specific to subclasses (Supplementary Table 5, [<u>Code</u>]), and for the cluster shown in Figure 1d, L6 CT, 324 isoforms belonging to 286 genes exhibiting isoform shifts in clusters (Supplementary Table 6, [<u>Code</u>]). Together, these form an isoform atlas for the MOp.

**Figure 2:**
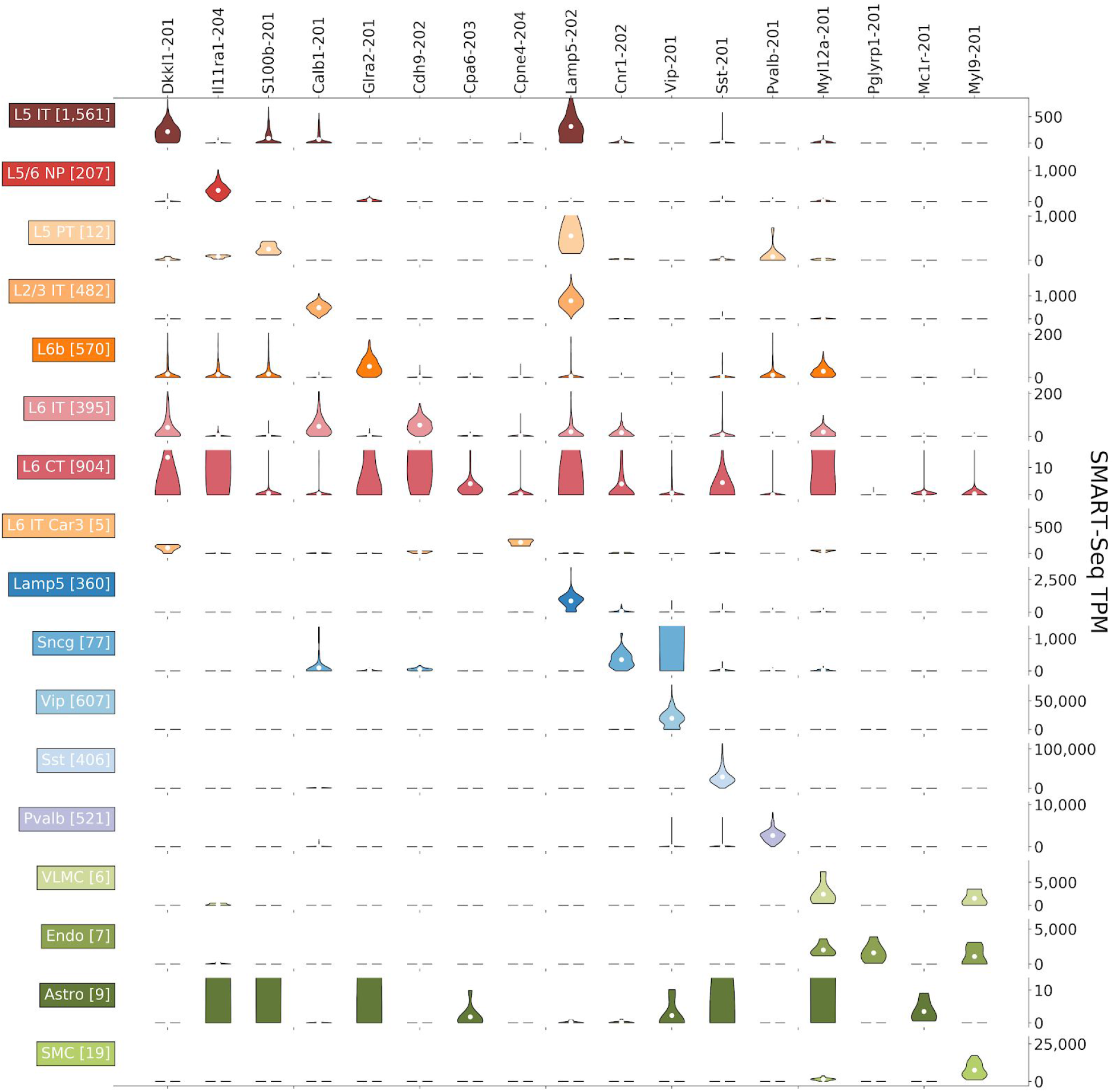
Isoform atlas. A sample from an isoform atlas displaying isoform markers differential with respect to subclasses. Each row corresponds to one subclass, and each column corresponds to one isoform. SMART-Seq isoform abundance estimates are in TPM units, and each column is scaled so that the maximum TPM is 4 times the mean of the isoform specific for that row’s cluster. [<u>Code</u>]

### Spatial isoform specificity

While spatial single-cell RNA-seq methods are not currently well-suited to directly probing isoforms of genes due to the number and lengths of probes required, spatial analysis at the gene-level can be refined to yield isoform-level results by extrapolating SMART-Seq isoform quantifications.

Figure 3a,b shows an example of a gene, Pvalb, where the SMART-Seq quantifications reveal that of the two isoforms of the gene, only one, Pvalb-201, is expressed. Moreover it can be seen to be specific to the Pvalb cell subclass (Figure 2). In an examination of MERFISH spatial single-cell RNA-seq, derived from 64 slices from the MOp region (Extended Data Fig. 7a), the Pvalb subclass, of which Pvalb is a marker, can be seen to be dispersed throughout the motor cortex spanning all layers (Extended Data Fig. 7b). While the MERFISH probes only measure abundance of Pvalb at the gene level (Figure 3c), extrapolation from the SMART-Seq quantifications can be used to refine the MERFISH result to reveal the spatial expression pattern of the Pvalb-201 isoform. This extrapolation can be done systematically. To build a spatial isoform atlas of the MOp, we identified differentially expressed genes from the MERFISH data (Supplementary Table 7, [<u>Code</u>]) and for each of them checked whether there were SMART-seq isoform markers (from Supplementary Table 5, [<u>Code</u>]). An example of the result is shown in Extended Data Fig. 8a, which displays one gene for each cluster, together with the isoform label specific to that cluster.

**Figure 3:**
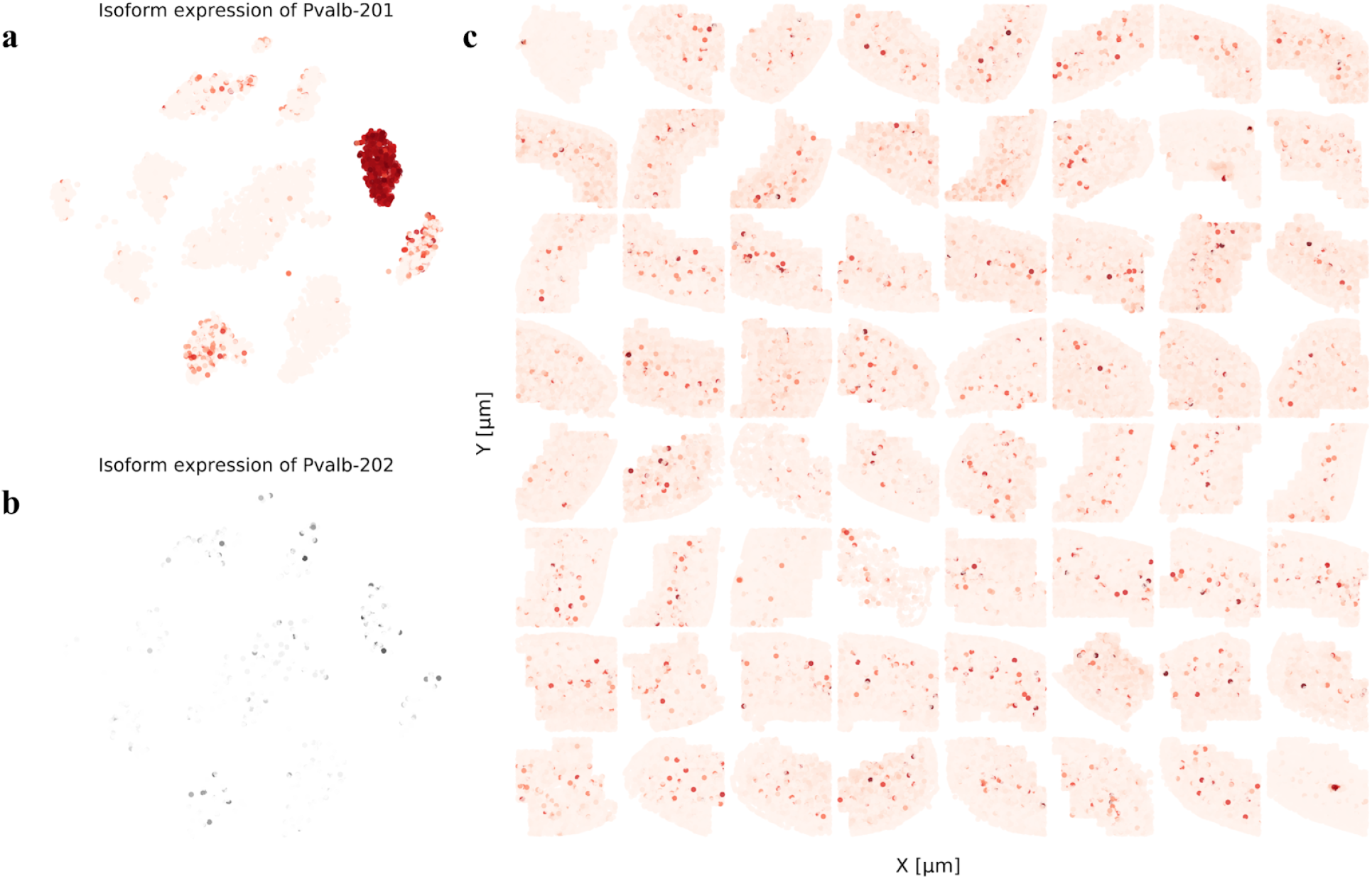
Spatial extrapolation of isoform expression. a) Spatial expression of the Pvalb-201 isoform across 64 slices from the MOp, as extrapolated from probes for the Pvalb gene. b) Expression of the Pvalb-201 isoform and c) expression of the Pvalb-202 isoform. [<u>Code a</u>. <u>Code b/c</u>]

While direct measurement of isoform abundance may be possible with spatial RNA-seq technologies such as SEQFISH^25^ or MERFISH^26^, such resolution would require dozens of probes to be assayed per gene (Extended Data Fig. 8b), each of which is typically tens of base-pairs in length. Thus, while isoforms can in theory be detected in cases where they contain large stretches of unique sequence, the technology is prohibitive for assaying most isoforms, making the extrapolation procedure described here of practical relevance.

### Splicing markers

Isoform quantification of RNA-seq can be used to distinguish shifts in expression between transcripts that share transcriptional start sites, and shifts due to the use of distinct transcription start sites. Investigating such differences can, in principle, shed light on transcriptional versus post-transcriptional regulation of detected isoform shifts^27,28^. Figure 4a shows an example of a gene, Rtn1, in which both GABAergic and glutamatergic classes exhibit similar preferential expression of transcripts at a specific start site (Figure 4b,c). However when examining the expression profile for the two isoforms within the highly expressed transcription start site, we observe that the glutamatergic class exhibits preferential expression of Rtn1-201, previously shown to be expressed in grey matter^29^, whereas the GABAergic class does not. We identified 29 isoforms that are preferentially expressed in either GABAergic, glutamatergic and Non-Neuronal classes, even when the expression of isoforms grouped by the same transcription start sites is constant among them (Supplementary Table 8, [<u>Code</u>]). Such cases are likely instances where the isoform shifts between cell types are a result of differential splicing, i.e. the result of a post-transcriptional program.

**Figure 4:**
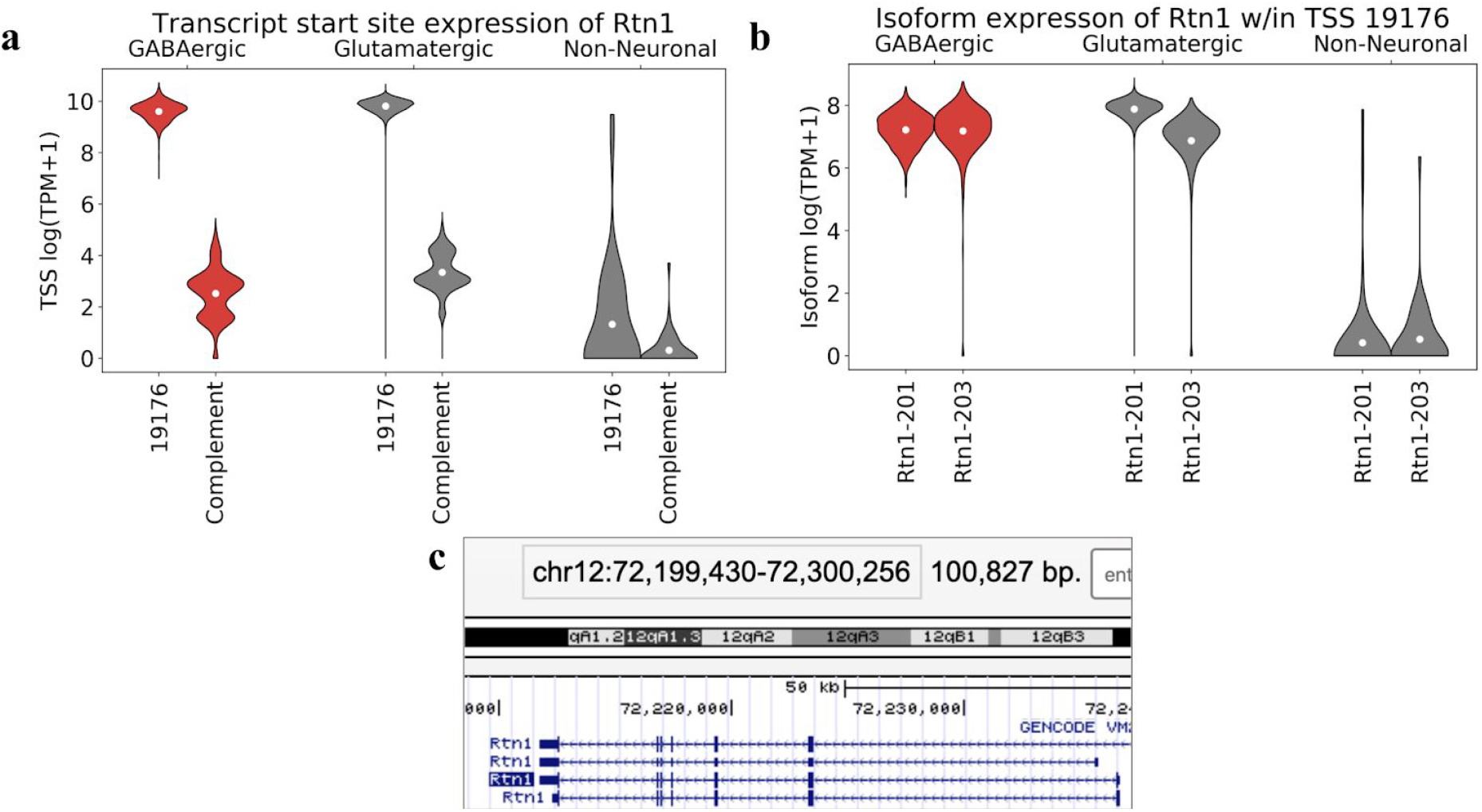
Isoform shifts reflecting splicing changes. a) Expression patterns of groups of transcript sharing the same TSS from the reticulon 1 (Rtn1) gene. b) Expression patterns of isoforms within TSS groups from the Rtn1 gene. c) The four isoforms of the Rtn1 gene. The 3rd and 4th isoforms from the top have the same transcription start site at the 5’ end of the transcript. [<u>Code a/b</u>, <u>Source c</u>]

## Discussion

Comparisons of different scRNA-seq technologies have tended to focus on throughput, cost, and gene-level accuracy^30^. Our results shed some light on the latter: it has been previously shown that quantification of isoform abundance is necessary for gene-level estimates^31^, and we verified that this is the case for SMART-Seq data. We found many examples of both false positive (Extended Data Fig. 9a) and false negative (Extended Data Fig. 9b) gene marker predictions at the subclass level (Supplementary Table 9, [<u>Code</u>] and Supplementary Table 10, [<u>Code</u>]). This highlights the importance of isoform quantification of SMART-Seq data, even for gene-centric analysis. In terms of accuracy compared to other technologies, we found slightly better agreement between SMART-Seq and MERFISH, two very different technologies, than between 10xv3 and MERFISH, for cell types where SMART-Seq assayed more than a handful of cells (Extended Data Fig. 10). This suggests that perhaps the “full-length” measurements of MERFISH and SMART-Seq confer an advantage over 3’-end based quantification, although more investigation is warranted.

While cost, throughput, and accuracy are important, we found the resolution of different technologies to be more fundamental in assessing their comparative advantages. Our results confirm that 10xv3 3’-end technology is better at detecting small cell populations than SMART-Seq (Extended Data Fig. 2a) due to the higher number of cells assayed. However SMART-seq has other advantages. The higher sensitivity of SMART-Seq leads to the identification of some cell types not detected by 10xv3, and we find that SMART-Seq’s full-length capture facilitates the identification of cell markers that cannot be detected from gene expression estimates produced with 3’-end methods (Figure 1, Extended Data Figs. 2b,c). Similarly, while spatial single-cell RNA-seq methods are, for the most part, currently limited to gene detection, refinement of expression signatures is possible by extrapolating SMART-Seq isoform quantification. In addition, isoform expression may result from transcriptional and post-transcriptional regulation within specific cell types that would be otherwise masked at the gene level.Thus, SMART-Seq is an important and powerful assay that can provide unique information not accessible with gene-centric technologies. Moreover, the recent development of a SMART-Seq protocol (SMART-Seq3), which can produce full-length and 5’-end reads simultaneously from single-cells^32^ may further improve the ability to register SMART-Seq cells with cells assayed using other technologies, thereby further increasing the possibilities for combining SMART-Seq with our scRNA-seq data. This perspective argues for using SMART-Seq data beyond naïve integration with other modalities at the gene level. The technology should be viewed as a complement, rather than competitor, to droplet or spatial single-cell RNA-seq. Our analyses suggest that a workflow consisting of droplet-based single-cell RNA-seq to identify cell types, then SMART-seq for isoform analysis, and finally spatial RNA-seq with a panel based on isoform-specific markers identified by SMART-seq, would effectively leverage different technologies’ strengths.

Our approach should be useful not only prospectively, but also for studies such as Kim et al. 2019 ^25^ which collected data using multiple scRNA-seq technologies but focused primarily on cell type identification using gene-level analyses. Our quantification of 6,160 SMART-Seq cells with kallisto^23^ shows that it is straightforward to produce accurate, reproducible quantifications efficiently for any SMART-Seq dataset. Moreover, our analysis, fully reproducible via python notebooks, can be used for a comprehensive analysis of any 10x Genomics Chromium, SMART-Seq and spatial scRNA-seq data. This should be useful not only for biological discovery of isoform functions related to cell types, but also as a form of validation of the clusters, since isoform markers are unlikely to be discovered within a cluster of cells by chance. Similarly, isoform markers can validate isoform quantification, because it is unlikely by chance that highly expressed isoforms, if produced by error, would cluster in a single group of cells.

The next step after assembling a single-cell isoform atlas is to probe the functional significance of cell type isoform specificity. Recently developed experimental methods for this purpose, e.g. isoforms screens^33^, are a promising direction and will be key to understanding the significance of the vast isoform diversity in the brain^34^.

## Supplementary Tables

Supplementary Table 1: Class level differential analysis with constant gene

Supplementary Table 2: Subclass level differential analysis with constant gene

Supplementary Table 3: Cluster level differential analysis with constant gene

Supplementary Table 4: Class level differential analysis with non-constant gene

Supplementary Table 5: Subclass level differential analysis with non-constant gene

Supplementary Table 6: Cluster level differential analysis with non-constant gene

Supplementary Table 7: Merfish gene level differential analysis

Supplementary Table 8: TSS level differential analysis

Supplementary Table 9: Naïve quantification gene level differential analysis

Supplementary Table 10: Valid quantification gene level differential analysis

## Methods

All of the results and figures in the paper are reproducible starting with the raw reads using scripts and code downloadable from https://github.com/pachterlab/BYVSTZP_2020. The repository makes the method choices completely transparent, including all parameters and thresholds used.

### Tissue collection and isolation of cells

Mouse breeding and husbandry: All procedures were carried out in accordance with Institutional Animal Care and Use Committee protocols at the Allen Institute for Brain Science. Mice were provided food and water ad libitum and were maintained on a regular 12-h day/night cycle at no more than five adult animals per cage. For this study, we enriched for neurons by using Snap25-IRES2-Cre mice^35^ (MGI:J:220523) crossed to Ai14^36^ (MGI: J:220523), which were maintained on the C57BL/6J background (RRID:IMSR_JAX:000664). Animals were euthanized at 53-59 days of postnatal age. Tissue was collected from both males and females (scRNA SMART, scRNA 10x v3).

Single-cell isolation: We isolated single cells by adapting previously described procedures^11,21^. The brain was dissected, submerged in ACSF^21^, embedded in 2% agarose, and sliced into 250-μm (SMART-Seq) or 350-μm (10x Genomics) coronal sections on a compresstome (Precisionary Instruments). The Allen Mouse Brain Common Coordinate Framework version 3 (CCFv3, RRID:SCR_002978)^37^ ontology was used to define MOp for dissections.

For SMART-Seq, MOp was microdissected from the slices and dissociated into single cells with 1 mg/ml pronase (Sigma P6911-1G) and processed as previously described^5^. For 10x Genomics, tissue pieces were digested with 30 U/ml papain (Worthington PAP2) in ACSF for 30 mins at 30 °C. Enzymatic digestion was quenched by exchanging the papain solution three times with quenching buffer (ACSF with 1% FBS and 0.2% BSA). The tissue pieces in the quenching buffer were triturated through a fire-polished pipette with 600-μm diameter opening approximately 20 times. The solution was allowed to settle and supernatant containing single cells was transferred to a new tube. Fresh quenching buffer was added to the settled tissue pieces, and trituration and supernatant transfer were repeated using 300-μm and 150-μm fire polished pipettes. The single cell suspension was passed through a 70-μm filter into a 15-ml conical tube with 500 ul of high BSA buffer (ACSF with 1% FBS and 1% BSA) at the bottom to help cushion the cells during centrifugation at 100xg in a swinging bucket centrifuge for 10 minutes. The supernatant was discarded, and the cell pellet was resuspended in quenching buffer.

All cells were collected by fluorescence-activated cell sorting (FACS, BD Aria II, RRID: SCR_018091) using a 130-μm nozzle. Cells were prepared for sorting by passing the suspension through a 70-μm filter and adding DAPI (to the final concentration of 2 ng/ml). Sorting strategy was as previously described^21^, with most cells collected using the tdTomato-positive label. For SMART-Seq, single cells were sorted into individual wells of 8-well PCR strips containing lysis buffer from the SMART-Seq v4 Ultra Low Input RNA Kit for Sequencing (Takara 634894) with RNase inhibitor (0.17 U/μl), immediately frozen on dry ice, and stored at −80 °C. For 10x Genomics, 30,000 cells were sorted within 10 minutes into a tube containing 500 μl of quenching buffer. Each aliquot of 30,000 sorted cells was gently layered on top of 200 μl of high BSA buffer and immediately centrifuged at 230xg for 10 minutes in a swinging bucket centrifuge. Supernatant was removed and 35 μl of buffer was left behind, in which the cell pellet was resuspended. The cell concentration was quantified, and immediately loaded onto the 10x Genomics Chromium controller.

### Genomic library preparation and sequencing

For SMART-Seq library preparation, we performed the procedures with positive and negative controls as previously described^21^. The SMART-Seq v4 (SSv4) Ultra Low Input RNA Kit for Sequencing (Takara Cat# 634894) was used to reverse transcribe poly(A) RNA and amplify full-length cDNA. Samples were amplified for 18 cycles in 8-well strips, in sets of 12-24 strips at a time. All samples proceeded through Nextera XT DNA Library Preparation (Illumina Cat# FC-131-1096) using Nextera XT Index Kit V2 (Illumina Cat# FC-131-2001) and a custom index set (Integrated DNA Technologies). Nextera XT DNA Library prep was performed according to manufacturer’s instructions, with a modification to reduce the volumes of all reagents and cDNA input to 0.4x or 0.5x of the original protocol.

For 10x v3 library preparation, we used the Chromium Single Cell 3’ Reagent Kit v3 (10x Genomics Cat# 1000075). We followed the manufacturer’s instructions for cell capture, barcoding, reverse transcription, cDNA amplification, and library construction. We targeted sequencing depth of 120,000 reads per cell.

Sequencing of SMART-Seq v4 libraries was performed as described previously^21^. Briefly, libraries were sequenced on an Illumina HiSeq2500 platform (paired-end with read lengths of 50 bp). 10x v3 libraries were sequenced on Illumina NovaSeq 6000 (RRID:SCR_016387).

### Pre-processing single-cell RNA-seq data

The 6,295 SMART-Seq cells were processed using kallisto with the ‘kallisto pseudo’ command^23^. The 94,162 10x Genomics v3 cells were pre-processed with kallisto and bustools^38^. Gene count matrices were made by using the --genecounts flag and TCC matrices were made by omitting it. The mouse transcriptome reference used was GRCm38.p3 (mm10) RefSeq annotation gff file retrieved from NCBI on 18 January 2016 (https://www.ncbi.nlm.nih.gov/genome/annotation_euk/all/), for consistency with the reference used by the BICCN consortium^17^.

The GTF and the GRCm38 genome fasta [<u>Source to both</u>], provided by the consortium, were used to create a transcriptome fasta, transcripts to genes map [<u>Source</u>], and kallisto index using kb ref -i index.idx, -g t2g.txt -f1 transcriptome.fa genome.fa genes.gtf.

Isoform and gene count matrices were generated for the Smart-seq2 data using the kallisto pseudo command. Cluster assignments were associated with cells using cluster labels generated by the BICCN consortium^17^. The labels are organized in a hierarchy of three levels: classes, subclasses and clusters. The cluster labels for the cells can be downloaded from https://github.com/pachterlab/BYVSTZP_2020.

### Normalization and filtering of SMART-Seq data

Isoform counts were first divided by the length of transcript to obtain abundance estimates proportional to molecule copy numbers. We then removed isoforms that had fewer than one count and that were in fewer than one cell. We also removed genes and their corresponding isoforms that had a dispersion of less than 0.001.

To generate the cell by gene matrix we summed the isoforms that correspond to the same gene. Cells with less than 250 gene counts and with greater than 10% mitochondrial content were removed. Cells were normalized to transcripts per million (TPM) by dividing the counts in each cell by the sum of the counts for that cell, then multiplying by 1,000,000. The count matrices were then transformed with log1p and the columns scaled to unit variance and zero mean. The resulting gene and isoform matrix contained 6,160 cells and 19,190 genes corresponding to 69,172 isoforms.

Highly variable isoforms and genes were identified by first computing the dispersion for each feature, and then binning all of the features into 20 bins. The dispersion for each feature was normalized by subtracting the mean dispersion and dividing by the variance of the dispersions within each bin. Then the top 5000 features were retained based on the normalized dispersion. This was computed by using the scanpy.pp.highly_variable_genes with n_top_genes = 5000, flavor=seurat, and n_bins=20^39^.

### Normalization and filtering of 10xv3 data

Isoform counts were first divided by the length of transcript to obtain abundance estimates proportional to molecule copy numbers. We then removed isoforms that had fewer than one count and that were in fewer than one cell. We also removed genes and their corresponding isoforms that had a dispersion of less than 1.

To generate the cell by gene matrix we summed the isoforms that correspond to the same gene. Cells with less than 250 gene counts and with greater than 21.5% mitochondrial content were removed. Cells were normalized to counts per million (CPM) by dividing the counts in each cell by the sum of the counts for that cell, then multiplying by 1,000,000. The count matrices were then transformed with log1p and the columns scaled to unit variance and zero mean. The resulting gene matrix contained 90,031 cells and 24,709 genes.

Highly variable isoforms and genes were identified by first computing the dispersion for each feature, and then binning all of the features into 20 bins. The dispersion for each feature was normalized by subtracting the mean dispersion and dividing by the variance of the dispersions within each bin. Then the top 5000 features were retained based on the normalized dispersion. This was computed by using the scanpy.pp.highly_variable_genes with n_top_genes = 5000, flavor=seurat, and n_bins=20^39^.

### Dimensionality reduction and visualization

Neighborhood component analysis^20^ (NCA) was performed on the full scaled log(TPM +1) matrix using the subcluster labels, to ten components. t-distributed stochastic neighbor embedding (t-SNE)^18^ was then performed on the 10 NCA components. t-SNE was computed using sklearn.manifold. t-SNE with default parameters and random state 42. Similarly uniform manifold approximation was performed on the 10 NCA components and the 50 truncated SVD components. Uniform Manifold Approximation and Projection (UMAP)^19^ was computed with the umap package with default parameters.

To ensure that NCA was not overfitting cells to their corresponding subclasses, we randomly permuted all of the subclasses labels and reran the NCA to t-SNE dimensionality reduction method. We observed uniform mixing of the permuted subclass labels, indicating that NCA was not overfitting the cells to their corresponding subclasses.

For the Louvain clustering displayed in Extended Data Fig. 5c, truncated SVD was performed on the 5000 top highly variable features and the first 50 SVD components were retained for the clustering. The random seed for all sklearn functions was 42, and default parameters were used for scanpy.pp.neighbors and scanpy.tl.louvain. The L5 IT subclass contains seven clusters in the SMART-Seq data, four clusters in the 10xv3 data, and three clusters in the MERFISH data.

### Measuring number of isoforms per gene

We parsed the transcripts to genes map, grouping together transcripts that had the same end site that were in the same gene. We then counted the number of these end site sets within a gene and plotted them against the number of isoforms within that gene.

### Cross-technology cluster correlation

The correlation between 10xv3-Smart-seq, 10xv3-MERFISH, and Smart-seq-MERFISH, was performed at the gene level and between cells grouped by subclasses for all three pairs of technologies, and at the isoform level and between cells grouped by cluster for only the 10xv3 and SMART-Seq. For each pair we started with two raw matrices and restricted to the set of genes/isoforms common to the two. Then we normalized the counts for each matrix per cell to one million, log1p transformed the entire matrix, and scaled the features to zero mean and unit variance. Within each cluster we restricted the features to those present in at least 50% of the cells. We then found the mean cell within the respective clusters in the two matrices, and computed the Pearson correlation between them. These methods were implemented for Extended Data Figs. 4--6.

### Isoform atlas

For each level of clustering: class, subclass, cluster, we performed a t-test for each gene/isoform between the cluster and its complement, on the log1p counts. To identify isoform enrichment that is masked at a gene level analysis, we looked for isoforms that were upregulated by checking that the gene containing that isoform was not significantly expressed in that cluster, relative to the complement of that cluster. Isoforms that were expressed in less than 90% of the cells in that cluster were ignored. All t-tests used a significance level of 0.01 and all p-values were corrected for multiple testing using Bonferroni correction.

### MERFISH isoform extrapolation

First we identified the genes that mark the specific subclass within the MERFISH data. The Pvalb gene is a marker for the Pvalb subclass. Then we performed differential analysis on the SMART-Seq data at the isoform level on the subclasses to identify the isoforms that mark each of the SMART-Seq subclasses. Only one of the two isoforms for Pvalb marked the Pvalb cluster. This allowed us to extrapolate the fact that the specific Pvalb isoform is being detected in the MERFISH data.

Additionally, we identified all of the genes that mark the specific subclasses in the MERFISH data through differential analysis and checked if their underlying isoforms were also differentially expressed. We then noted which isoforms were differentially expressed for the spatial isoform atlas.

### Grouping transcripts by start site

Using the transcripts to genes map and the filtered isoform matrix generated before, we grouped isoforms by their transcription start site into TSS classes and summed the raw counts for the isoforms within each TSS class to create a *cellx TSS* matrix. Differential analysis was then performed in exactly the same way as above. For each cluster and each TSS/isoform, a t-test was performed between the cells in that cluster and the cells in the complement of that cluster. All statistical tests used a significance level of 0.01 and all p-values were corrected for multiple testing using Bonferroni correction.

### Comparison of naïve and valid quantification

Naïve gene count matrices were constructed from the SMART-Seq data by summing the counts corresponding to a single gene. Valid gene count matrices were made with SMART-Seq by first dividing isoform abundances by the length of their transcripts, and then summing the abundances of isoforms by gene. Differential analysis was performed independently on these two gene count matrices. and the resultant differential genes were compared. Subsequently false-positive and false-negative detections of differentially abundant genes were reported.

## Software versions

<u>Anndata 0.7.1</u>

<u>bustools 0.39.4</u>

<u>awk (GNU awk) 4.1.4</u>

<u>grep (GNU grep) 3.1</u>

<u>kallisto 0.46.1</u>

<u>kb_python 0.24.4</u>

<u>Matplotlib 3.0.3</u>

<u>Numpy 1.18.1</u>

<u>Pandas 0.25.3</u>

<u>Scanpy 1.4.5.post3</u>

<u>Scipy 1.4.1</u>

<u>sed (GNU sed) 4.4</u>

<u>sklearn 0.22.1</u>

<u>tar (GNU tar) 1.29</u>

<u>umap 0.3.10</u>

## Data and Software Availability

The software used to generate the results and figures of the paper is available at https://github.com/pachterlab/BYVSTZP_2020. The single-cell RNA-seq data used in this study was generated as part of the BICCN consortium^17^. The 10xv3 and SMART-Seq data can be downloaded from http://data.nemoarchive.org/biccn/lab/zeng/transcriptome/scell/. The MERFISH data was generated as part of the BICCN consortium but is not currently available due to restrictions by Xiaowei Zhuang; requests for the data can be made directly to her by emailing.

## Acknowledgments

We thank members of the BICCN consortium, especially the “Mini-MOp” analysis group, for helpful conversations related to transcriptome analysis of the primary motor cortex. Thanks to Nadezda Volovich, Vasilis Ntranos, and Páll Melsted, for help with a preliminary quantification of the SMART-Seq data. This work was funded by the NIH Brain Initiative via grant U19MH114930 to HZ and LP).

## Author Contributions

ASB and LP conceived the study. ASB implemented the methods and produced the results and figures. ASB and LP analyzed the data and wrote the manuscript. ZY, CV, KS, BT and HZ produced the SMART-Seq and 10xv3 data.

