## Supplementary Figures for "Isoform cell type specificity in the mouse primary motor cortex"

**Note**: The captions of the Extended Data Figs. contain links to the code used to make the figures.

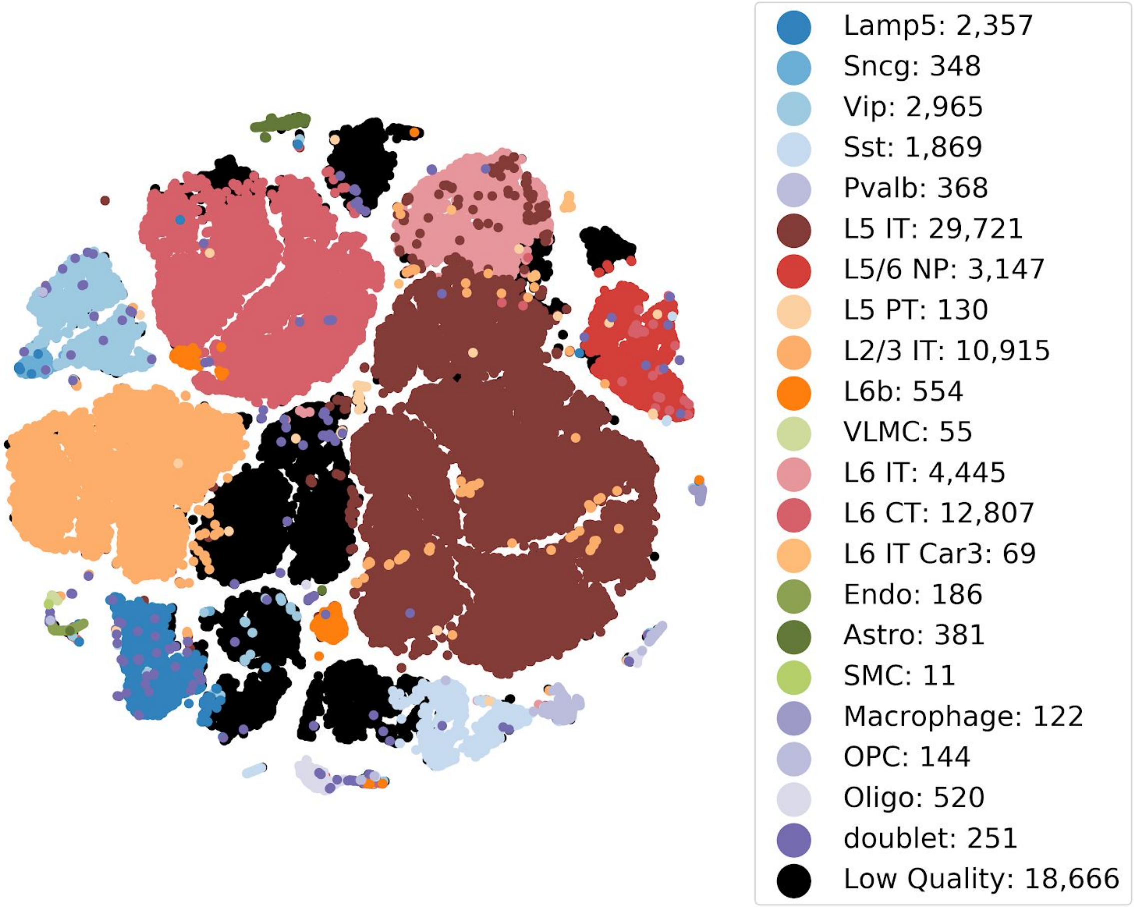


**Extended Data Figure 1:** t-distributed stochastic neighbor embedding of 90,031 10xv3 cells from the mouse primary motor cortex annotated with cell subclass assignments. The number of cells in each subclass is displayed next to the subclass label. [Code]

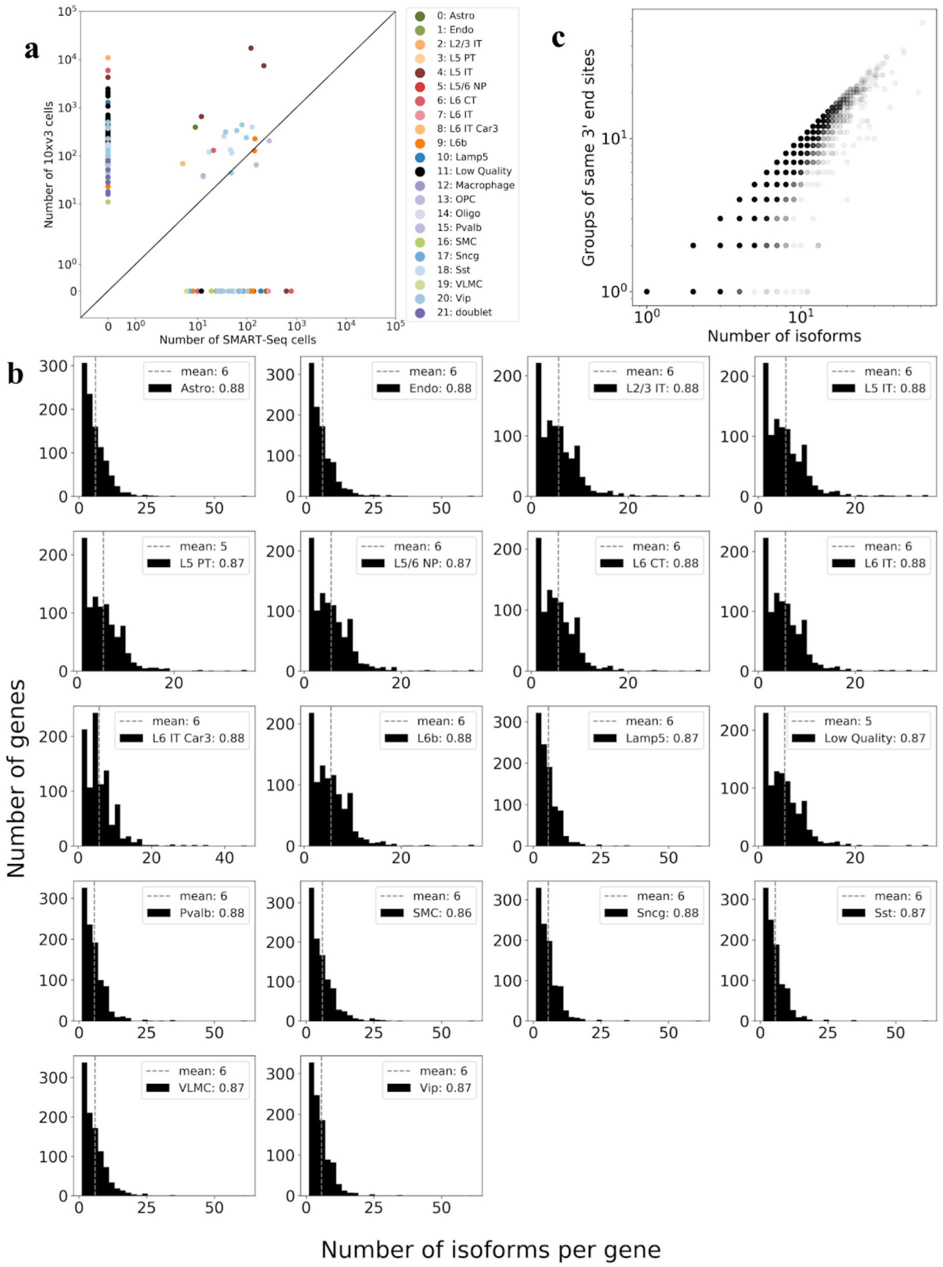

**Extended Data Figure 2:** a) Comparison of subclass identification for 10xv3 and SMART-Seq. Subclasses were identified separately, but with the same method for data from each of the technologies. 125 clusters with gene markers were identified in the 10xv3 data but not in the SMART-Seq data while 40 clusters with gene markers were identified in the SMART-Seq data and not the Chromium data. b) The distribution of the number of isoforms per gene within each of 18 subclasses, computed from the top 998 most highly expressed genes in the SMART-Seq dataset. The number associated with each class indicates the fraction of genes for which there are more than one isoform. c) Extent of isoform diversity in groups of transcripts sharing a 3' end. Each point displays the density of the number of groups (y-axis) containing a given number of isoforms (x-axis). Points along the line $y = x$ correspond to transcripts with unique 3' ends. [Code a, Code b, Code c]

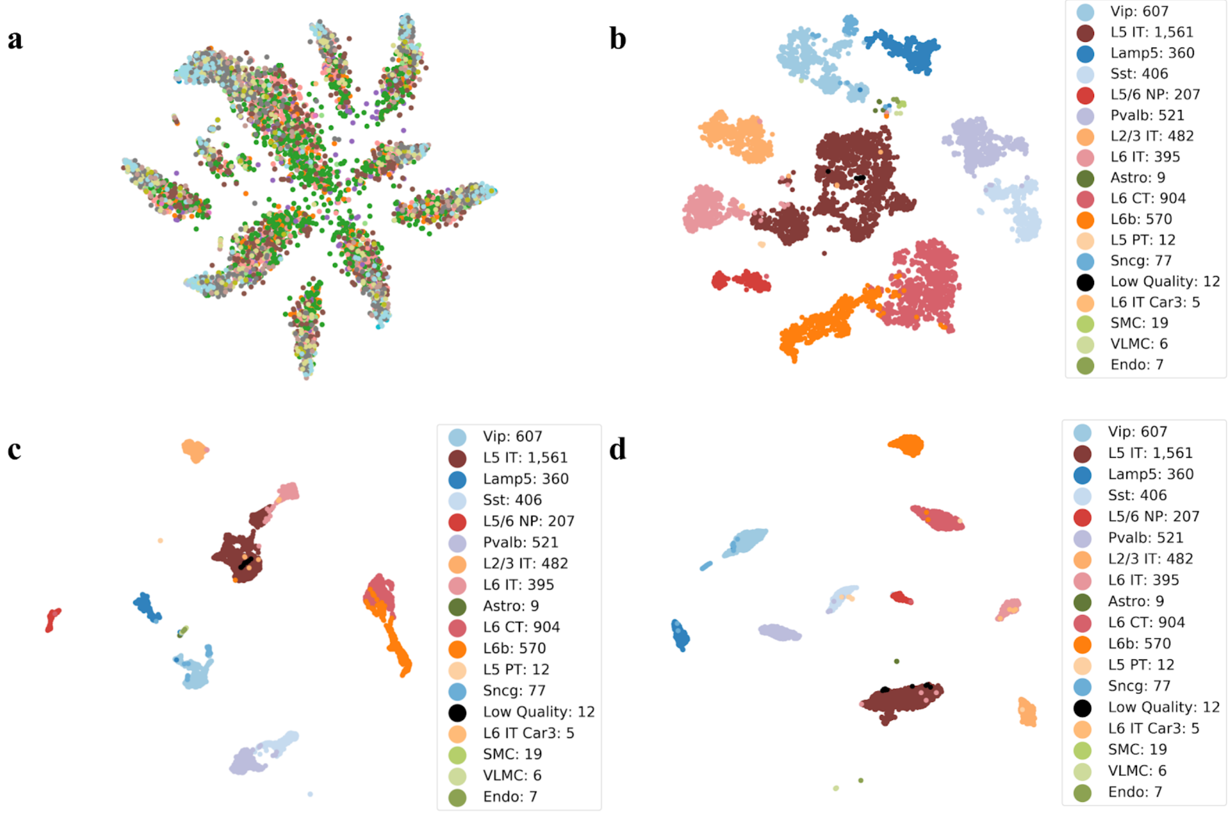


**Extended Data Figure 3:** a) Neighborhood component analysis on the scaled log1p normalized SMART-Seq gene matrix with the subclass labels permuted randomly to ten components, followed by t-SNE to dimension two. The lack of separation of cells by label demonstrated that the NCA procedure is not overfitting the data. b) Truncated SVD on the scaled log1p normalized SMART-Seq gene matrix to 50 components followed by t-SNE to two dimensions. The numbers next to each subclass label indicate the number of cells in the subclass. c) Truncated SVD on the scaled log1p normalized SMART-Seq gene matrix to 50 components followed by UMAP to dimension two. The numbers next to each subclass label indicate the number of cells in the subclass. d) Neighborhood component analysis on the scaled log1p normalized SMART-Seq gene matrix followed by UMAP to dimension two. The numbers next to each subclass label indicate the number of cells in the subclass. [Code a, Code b, Code c, Code d]

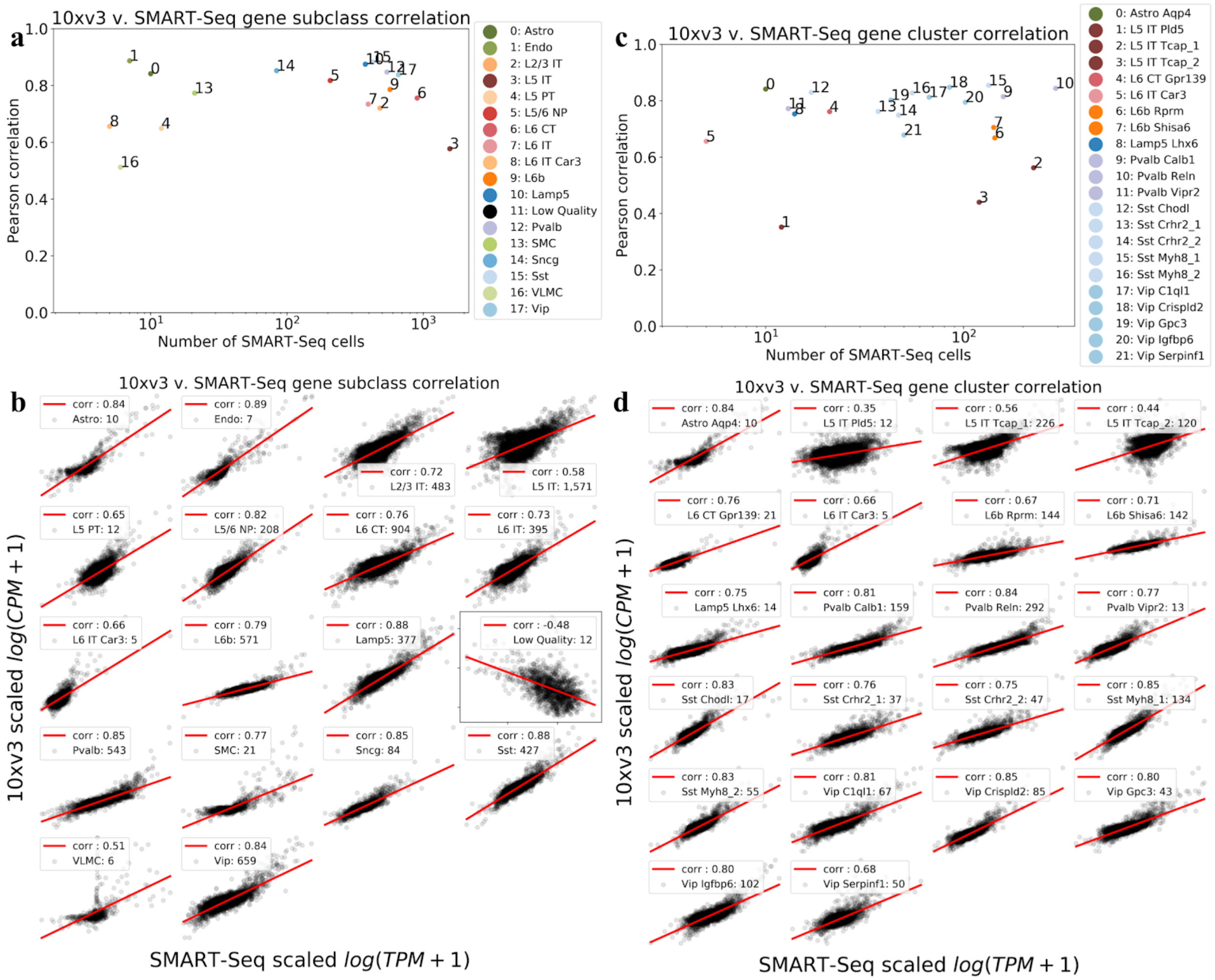


**Extended Data Figure 4:** a) Pearson correlation by subclass of the mean gene expression in 10xv3 and the mean gene expression in SMART-Seq, against the size of the subclass, for genes that are expressed in at least 50% of cells in that subclass. b) Scatter plot by subclass of the mean gene expression in 10xv3 vs the mean gene expression in SMART-Seq for genes that are expressed in at least 50% of cells in that subclass. The subclass sizes and Pearson correlation values are also reported. c) Pearson correlation by cluster of the mean gene expression in 10xv3 and the mean gene expression in SMART-Seq, against the size of the cluster, for genes that are expressed in at least 50% of cells in that cluster. d) Scatter plot by cluster of the mean gene expression in 10xv3 vs the mean gene expression in SMART-Seq for genes that are expressed in at least 50% of cells in that cluster. The cluster sizes and Pearson correlation values are also reported. [Code a, Code b, Code c, Code d]

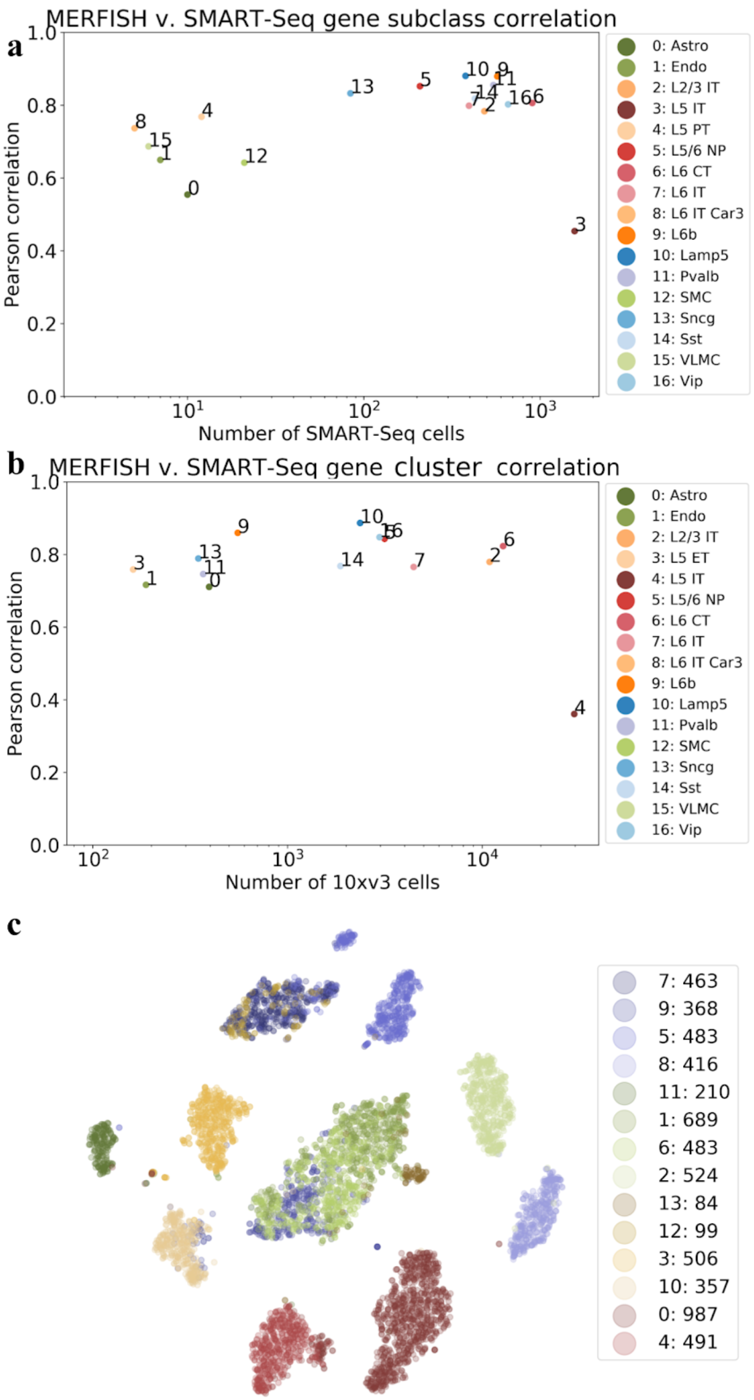


**Extended Data Figure 5:** a) Pearson correlation by subclass of the mean gene expression in MERFISH and the mean gene expression in SMART-Seq, against the size of the subclass, for genes that are expressed in at least 50% of cells in that subclass. b) Pearson correlation by subclass of the mean gene expression in MERFISH and the mean gene expression in 10xv3, against the size of the subclass, for genes that are expressed in at least 50% of cells in that subclass. c) Louvain clustering of the scaled log1p normalized gene count matrix. The central cluster, corresponding to the L5 IT cluster, is comprised of discordant subclasses. [Code a, Code b, Code c]

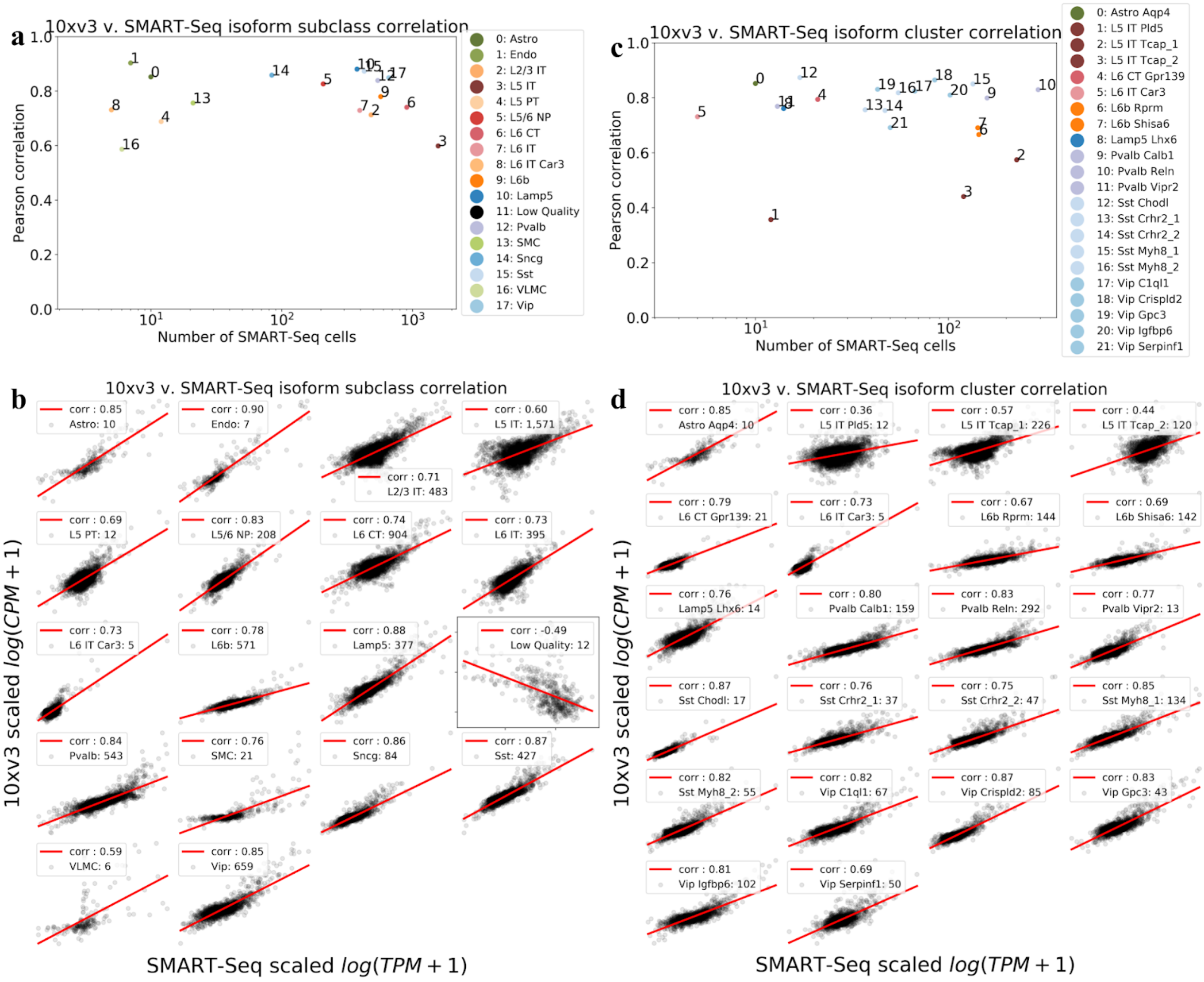


**Extended Data Figure 6:** a) Pearson correlation by subclass of the mean isoform expression in 10xv3 and the mean isoform expression in SMART-Seq, against the size of the subclass, for isoforms that are expressed in at least 50% of cells in that subclass. b) Scatter plot by subclass of the mean isoform expression in 10xv3 vs the mean isoform expression in SMART-Seq for isoforms that are expressed in at least 50% of cells in that subclass. The subclass sizes and Pearson correlation values are also reported. c) Pearson correlation by cluster of the mean isoform expression in 10xv3 and the mean isoform expression in SMART-Seq, against the size of the cluster, for isoforms that are expressed in at least 50% of cells in that cluster. d) Scatter plot by cluster of the mean isoform expression in 10xv3 vs the mean isoform expression in SMART-Seq for isoforms that are expressed in at least 50% of cells in that cluster. The cluster sizes and Pearson correlation values are also reported. [Code a, Code b, Code c, Code d]

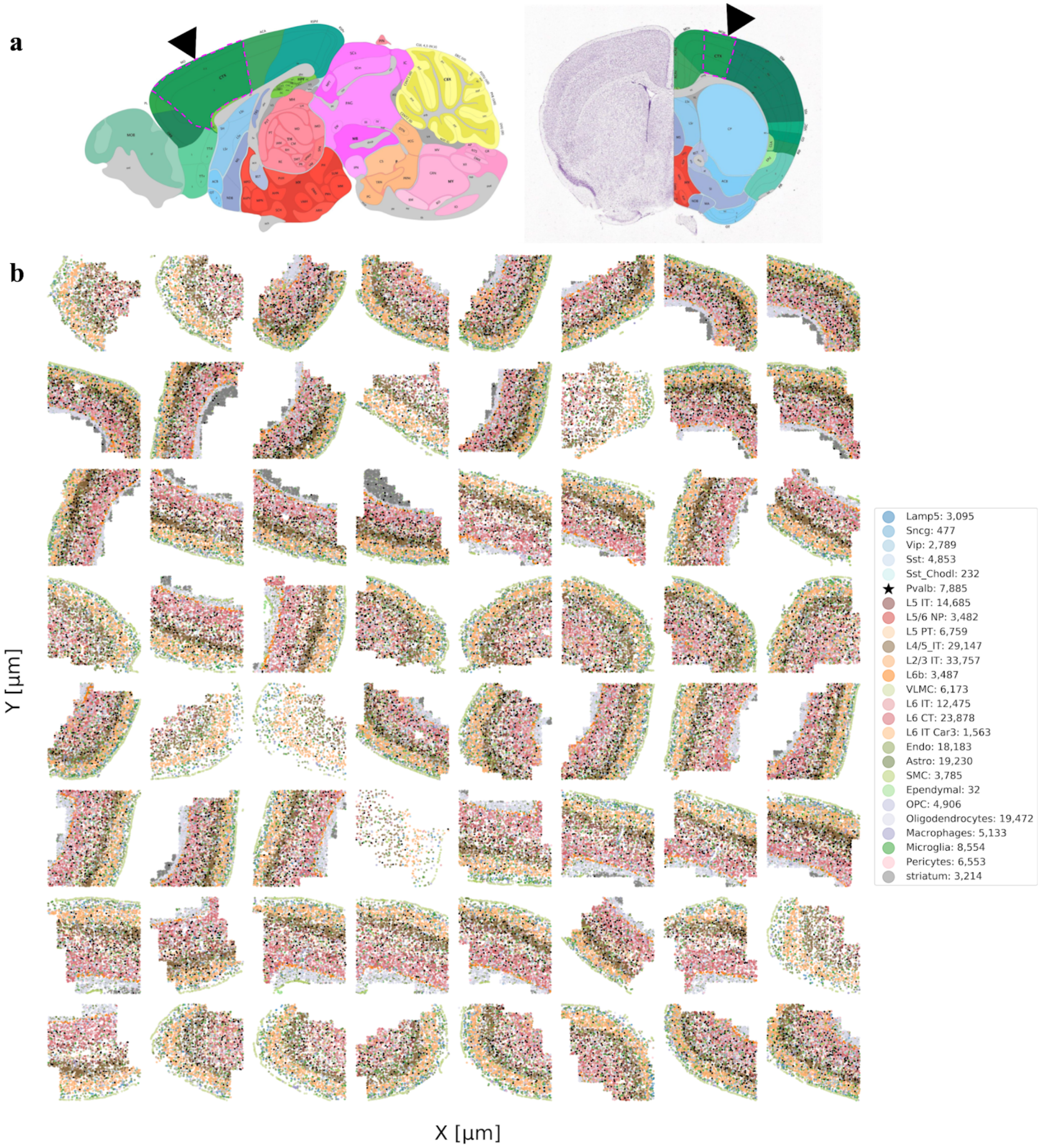


**Extended Data Figure 7:** a) The location of the mouse primary motor cortex, outlined in pink and pointed to by a black arrow. Shown are the sagittal view (left) and coronal view (right). b) Spatial location of cells in all subclasses across 64 slices from the MOp, the Pvalb cells are starred. [Source a, Code b]

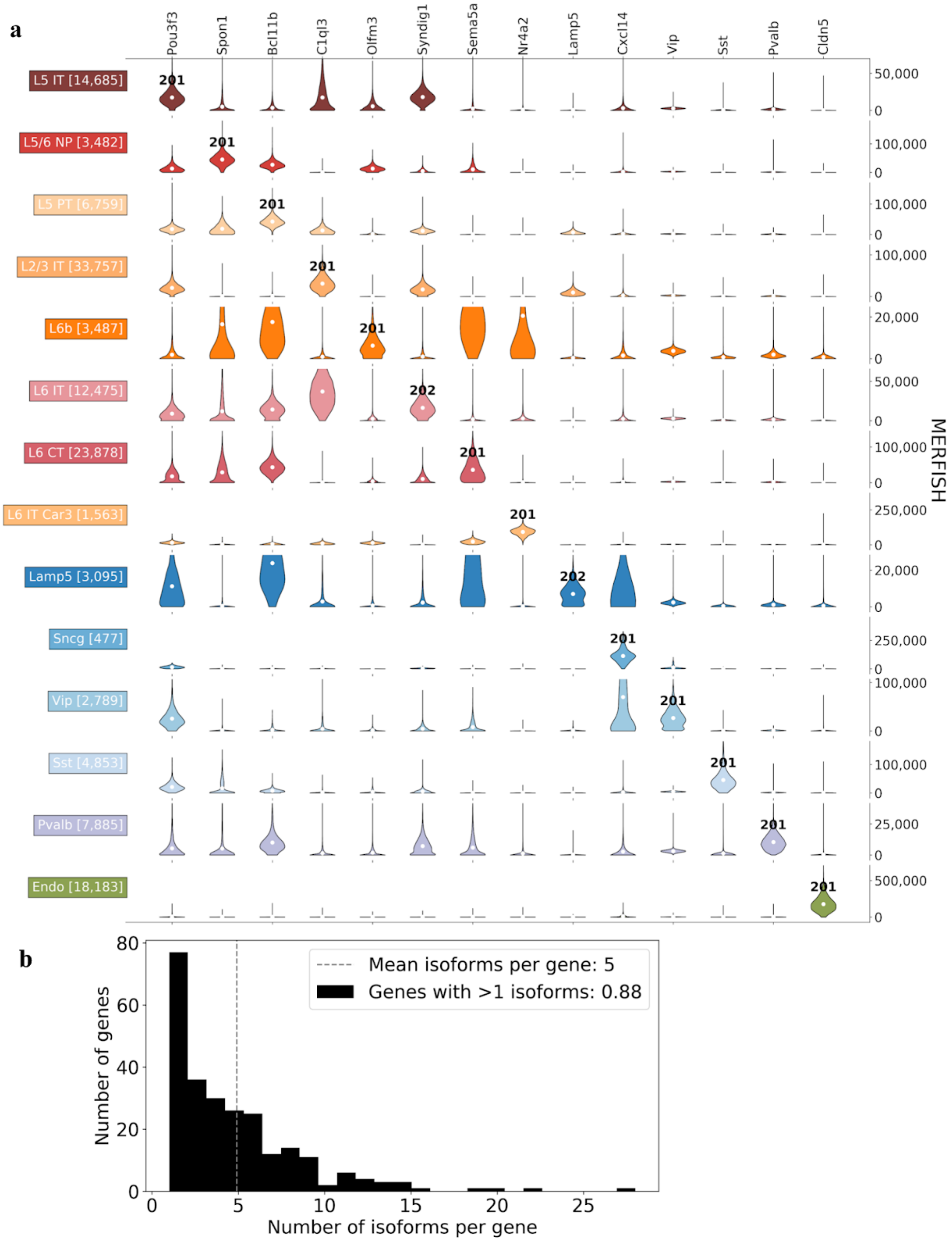


**Extended Data Figure 8:** a) Spatial isoform atlas for MERFISH data. Each column corresponds to a marker gene where there exists an underlying isoform that was differential in the SMART-Seq data, and which was also differential in the MERFISH data. The normalized gene expression values are plotted for each cluster-gene pair. The isoform that is differential for each cluster is displayed in bold on the corresponding violin plot. b) The distribution of the number of isoforms per gene for all of the 254 genes targeted by MERFISH. [Code a, Code b]

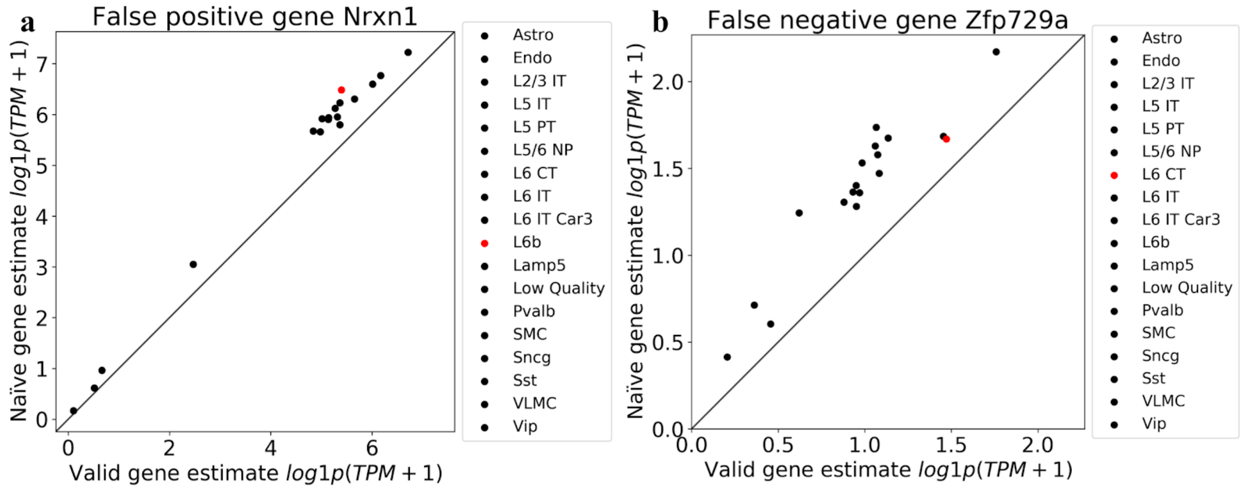


**Extended Data Figure 9:** Comparison of two quantifications approaches: "valid" gene quantification where isoform abundances are estimates, and then added up to obtain a gene abundance estimate, versus naive quantification in which read counts are collated by gene locus. a) In this case naïve quantification results in a false positive gene marker for the L6b subclass. b) In this case naïve quantification results in a false negative gene marker for the L6 CT subclass. [Code a, Code b]

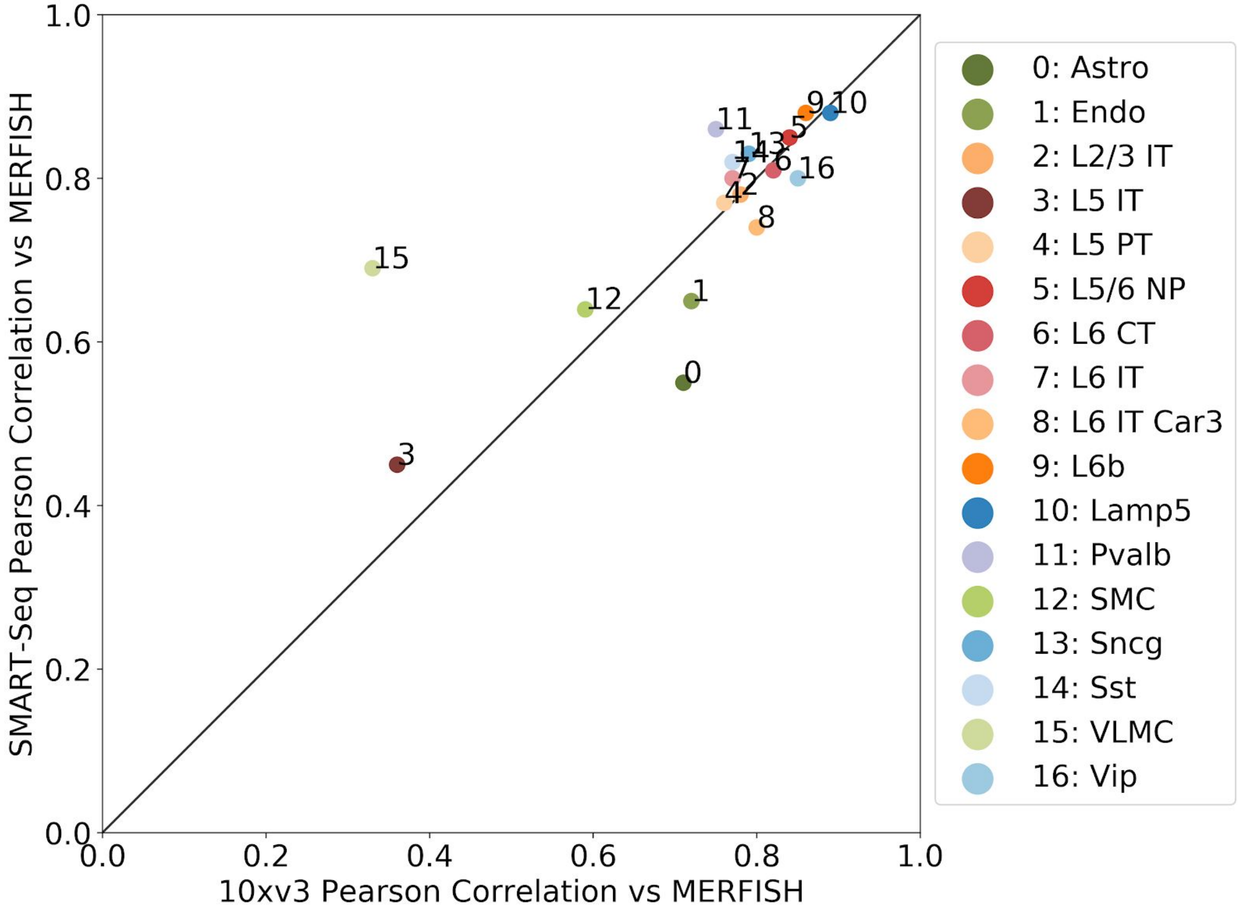


**Extended Data Figure 10:** Comparison of gene correlations by cell type between 10xv3 and MERFISH, and SMART-Seq and MERFISH computed using the 254 genes assayed in the MERFISH dataset. [Code]
